## Supporting information for "Sulfite preservatives effects on the mouth microbiome: changes in viability, diversity and composition of microbiota"

**Sulfite Exposure Protocol**:

To prepare samples, either sterile DI water (for controls), 1 M Na_2_SO_3_ (sodium sulfite treated), or 1M NaHSO_3_ (sodium bisulfite treated) was added to each saliva sample. For “time 0”, each treated sample was immediately centrifuged at 4,600 RPMs for 10 minutes, the supernatant discarded and the pellet brought up in 1X phosphate buffered saline (PBS). Simultaneous to placing time 0 treated and control samples in the centrifuge, the remaining portions of the prepared samples were incubated at 36°C for 30 minutes. After incubation, these samples were briefly centrifuged and mixed with a pipet before removing saliva to be centrifuged. Samples were centrifuged at 4,600 RPM for 10 minutes, the supernatant discarded and the pellet brought up in 1X PBS. All samples were frozen at -20℃ for less than 2 weeks or -80℃ for longer storage.

All control time zero samples were prepared, up until being centrifuged, prior to preparing treated samples in order to limit saliva exposure time to sulfites. For further caution in this regard, saliva was exposed to a given sulfite (NaHSO₃ or Na₂SO₃) in separate experiments. Noted time points, Time 0 and Time 30, are based on sulfite exposure time preceding the centrifuge process.

Challenges of working with saliva to evaluate the mouth microbiome have been noted in several studies. Factors to be considered include the constant turnover of saliva, changes in the environment due to ingestion of food and beverages, oral hygiene, age, smoking, and gender. In addition, the viscosity and heterogeneity of most samples makes it difficult to mix and pipette to a consistently high degree of accuracy. To reduce the pipetting errors each sample was initially diluted 1:10 with sterile water before aliquoting. Another possible source of error to be aware of, is in the pelleting and resuspension of cells in saliva after treatments. This was done to remove sulfites from cells and effectively stop the reaction. In some cases, cells did not form as solid of a pellet making it more difficult to remove supernatant without also removing some cells. We hypothesize that the differences we observed in pellet formation is due in part to differences in the chemical makeup of each saliva sample. Future studies may forgo the pelleting step and proceed with ATP test in saliva rather than PBS to avoid these errors, however that will present other problems with the exposure time and if samples are then to be used in sequencing they will likely need to be processed immediately.  We chose to use this method despite some variability observed in control samples used in each test as the difference was not significant. We also note that despite these known sources of error, the decrease in ATP activity observed in both preliminary work and in these experiments was significant and consistent.

**Evaluating ATP Data and Calculations:** The RLU (ATP activity) readings after the BacTiter-Glo™ Microbial Cell Viability Assay were examined further to allow data comparison. Each sample was tested in triplicate. If one of the samples reading indicated a 50% decrease or increase from the other two samples, they were omitted. Otherwise the replicates were averaged and background subtracted. The background was determined for each plate by placing controls of water and buffer reagent in a set of designated triplet wells. There were six sample dates from three individuals with sodium sulfite exposure, and two individuals with sodium bisulfite exposure whose results for ATP were omitted completely due to experimental error. This was less than 2% of samples tested and did not include any samples from individual F2. After reviewing the data, the control and treated values of ATP activity were compared and evaluated for percent change.

ATP activity in individual F2’s control samples vs lysozyme activity (initial rate) is presented in Figure S1. As also shown in Fig.3 which compared all 10 individuals’ samples, we did not observe any consistent correlation between ATP activity and lysozyme activity in these samples.

**Fig S1. F2 viable cells and lysozyme activity in control samples.** Average ATP activity (RLU) based on replicates of 3 for each sample of untreated control samples compared to the average Lysozyme activity (AFU) in the same samples of individual F2. Standard deviation for replicates is shown.
