## Supplementary material for "Sulfite preservatives effects on the mouth microbiome: changes in viability, diversity and composition of microbiota": S2

**Individual F2** Time 0

**[DNA]ng/ul** 10 minute exposure time

| | control | sodium bisulfite | |
|---|---|---|---|
| Sample | | | |
| 9A | 0.256815 | 3.333307 | 9B |
| 10A | 1.234404 | 1.179294 | 10B |
| 11A | 3.610662 | 0.405481 | 11B |
| 12A | 0.104199 | 0.891518 | 4B |
| 13A | 0.643877 | 2.00543 | 12B |
| 14A | 1.248576 | 0.389599 | 13B |
| 15A | 3.042318 | 2.563435 | 14B |
| 16A | 4.914455 | 1.559958 | 16B |
| Average | 1.881913 | 1.541003 | |

Average -18% drop in DNA concentration with sodium bisulf

p=0.34 0.336346 one tailed T-Test

| | Control | sodium sulfite | |
|---|---|---|---|
| 1A | 2.113692 | 0.016807 | 1B |
| 2A | 0.946527 | 0.804895 | 2B |
| 3A | 0.001195 | 0.002339 | 3B |
| 4A | 0.009235 | 0.004531 | 4B |
| 5A | 0.93988 | 0.660755 | 5B |
| 6A | 0.26809 | 0.137254 | 6B |
| 7A | 1.394653 | 0.913752 | 7B |
| 8A | 0.933279 | 0.384036 | 8B |
| Average | 0.825819 | 0.365546 | |

Average -56% drop in DNA concentration with sodium sulfite

p =0.051 one tailed T test

ite exposure
